# Testing the myth of humid versus dry cold: birds do not care

**DOI:** 10.1101/789792

**Authors:** Ryan S. O’Connor, François Vézina

## Abstract

At high air temperatures (T_a_) atmospheric water vapor can markedly influence an animal’s heat dissipation by reducing evaporative cooling capacities. At low T_a_ however, the impact of increased water vapor on heat exchange has sparsely been investigated, despite human populations at northern latitudes insisting on feeling colder on days with higher humidity (i.e., humid-cold). Here, we aimed to investigate the humid-cold perception by determining whether Black-capped Chickadees (*Poecile atricapillus*) exhibited greater energy expenditure through increased heat loss under humid-cold conditions relative to dry-cold. We measured resting metabolic rates (RMR) of captive chickadees (n=10) during exposure to two T_a_ treatments, below freezing (T_a_ ≈ −7°C) and above freezing (T_a_ ≈ 10°C), with either dry or saturated air. We found that T_a_ substantially impacted RMR, with RMR ≈1.5-fold greater at −7°C compared with 10°C. Conversely, humidity did not have a statistically significant impact on RMR at either T_a_. Our data thus suggest that humidity does not significantly influence an individual’s heat exchange with the environment at cold T_a_. This presumably reflects the fact that cold air can hold minimal amounts of water vapor and any possible influence on heat loss is negligible. Instead, we suggest that a greater contributor to the humid-cold myth is the frequency of overcast days during which direct solar radiation is blocked.

**Summary statement:** Black-capped chickadees significantly increased their resting metabolic rates at air temperatures below freezing relative to air temperatures above freezing, whereas humidity level did not significantly affect resting metabolic rates.

## INTRODUCTION

Heat loss to the environment is an ever-present challenge for endothermic animals living in cold regions. Consequently, a substantial portion of an endotherm’s daily energy budget in cold environments is dedicated towards thermoregulation (Raichlen et al., 2010; Burton et al., 2011). An individual’s energetic expenditure may be further influenced by climatic factors (e.g., wind and solar radiation) which can alter rates of heat loss and/or heat gain (Robinson et al., 1976; Hayes and Gessaman, 1980; Bakken et al., 1991; Walsberg and Wolf, 1995). Geiser and Drury (2003), for example, demonstrated that the energetic cost for Stripe-faced Dunnart’s (*Sminthopsis macroura*) to actively rewarm (i.e., shiver) from torpor was 6.3 times greater compared with individual’s that passively rewarmed under radiative heat. Conversely, in warm environments where heat dissipation is paramount, increasing humidity can considerably impact an organism’s capacity to evaporatively cool (Webster and King, 1987; Powers, 1992; Gerson et al., 2014).

Among human populations living at northern latitudes it is common to “feel colder” on days with higher relative humidity (RH; https://www.theglobeandmail.com/life/health-and-fitness/fitness/is-damp-cold-really-the-worst/article28346087/). However, this perception appears largely qualitative and few investigations have sought to quantify its validity. It is suggested that more humid air makes one feel colder relative to dry conditions because the water vapor increases the thermal conductivity of air resulting in a greater rate of heat transfer away from the body (Crow, 1988). Yet, at air temperatures (T_a_) below freezing, the amount of water vapor present in the air is minimal relative to conditions above freezing because most water will have condensed out. Consequently, at T_a_ < 0°C, the amount of water vapor present in the air to increase the conductance of heat away from the body is greatly reduced and should have little influence on heat loss. Thus, although the tenacious belief of humid-cold persists in northern latitudes where T_a_ regularly falls below freezing, a humid cooling effect should be more myth than reality.

In a recent study, Petit and Vézina (2014) investigated the thermal energetics of Black-capped Chickadees (*Poecile atricapillus*) wintering in a region where minimum T_a_ falls below freezing for 5-6 months of the year. Although Petit and Vézina (2014) found that T_a_ was the main driver for seasonal variation in mass and sex-independent summit metabolic rates (M_sum_), they also observed higher thermogenic capacity on days with higher absolute humidity. It is noteworthy, however, that this response was limited to days when T_a_ was above freezing and Petit and Vézina (2014) cautioned that the amount of variation in M_sum_ that was explained by absolute humidity was very small. Moreover, M_sum_ is an experimental measurement of maximal shivering heat production when exposed to helox gas, an experimental setup which creates extremely cold and dry environments (Rosenmann and Morrison, 1974). Summit metabolism is thus likely not a good indicator of heat loss due to humidity under conditions of less extreme cold.

In the present study, our main objective was to quantify the impact of humidity on heat loss in cold environments. We argue that if a cooling effect of humid air indeed exists it should increase heat production in an endothermic animal when T_a_ is, (i) above freezing, with more water vapor present in the air, and (ii) below the lower critical temperature of thermoneutrality (LCT; the temperature below which active heat production becomes necessary for maintaining a stable and elevated body temperature; Scholander et al., 1950). To determine the influence of humidity on heat loss, we compared resting metabolic heat production in Black-capped Chickadees exposed to dry and humid air at T_a_ below and above freezing. We predicted that at T_a_ < 0°C, the effect of humidity on heat loss would be negligible relative to dry conditions, leading to similar metabolic rates. Conversely, under conditions when T_a_ > 0°C, but below the LCT, we predicted metabolic rates would be higher in individuals exposed to humid air relative to those in a dry environment, a result expected if a humid-cooling effect existed.

## MATERIALS AND METHODS

### Experimental birds and design

This experiment was conducted between 20 February and 26 March 2016. Black-capped Chickadees (n = 10) used in this study were a subset of captive birds that were held in outdoor aviaries at the Université du Québec à Rimouski. Throughout the experiment, birds were provided food and water *ad libitum*. Food was removed one hour before metabolic measurements to ensure post-absorptive conditions.

Our experimental design consisted of measuring resting metabolic rates (RMR) under two T_a_ treatments, namely below freezing (set T_a_ = −10°C) and above freezing (set T_a_ = 10°C), and under two levels of humidity. Hence, all birds experienced the following conditions: (i) below freezing with dry air, (ii) below freezing with humid air, (iii) above freezing with dry air, and (iv) above freezing with humid air. The order in which each bird experienced a treatment was randomized to avoid potential habituation effects (Jacobs and McKechnie, 2014). We chose a set T_a_ of −10°C for the below freezing treatment because this corresponds to the mean minimal T_a_ experienced by our source population during January and February (Petit and Vézina, 2014). However, heat dissipated from a bird’s body increased the average T_a_ inside chambers and we thus report average T_a_ values in the results as opposed to the set T_a_. The second set T_a_ of 10°C was chosen because it is below the LCT of Black-capped Chickadees (Cooper and Swanson, 1994).

### Resting metabolic rate, air humidification, and experimental protocol

We used open, flow-through respirometry to measure water vapor pressure (WVP; measured in kilopascals, kPa) and oxygen consumption (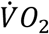; mL min^−1^). All RMR measurements were performed on two birds simultaneously and measured between 10:00 and 17:00 hrs. Before each experiment, we measured a bird’s body mass (M_b_) and body temperature (T_b_). Body temperature was measured in the cloaca using a type T thermocouple and an Omega Engineering thermocouple reader (model CNi3253, Norwalk, CT, USA).

Birds were placed individually in 1-L stainless steel metabolic chambers that were subsequently placed inside a modified household chest-freezer which acted as a temperature-controlled cabinet. Atmospheric air was scrubbed of CO_2_ and H_2_O using columns of soda lime and Drierite, respectively, connected in series. Once scrubbed, we split the air stream into 3 channels, representing a baseline and two chamber channels. Flow rates of incurrent air entering the metabolic chambers were held constant throughout an experiment at 500 mL min^−1^ using an Omega Engineering mass flow controller (model FMA5400A/FMA5500A, Norwalk, CT, USA) previously calibrated against a bubble-O-meter (Dublin, OH, USA). All tubing used in the system was Bev-a-line tubing.

To manipulate the water vapor content of incurrent air, we used 1-L Nalgene bottles, containing approximately 750 mL of water, as a bubbling device at room temperature (≈21°C). Dry air exiting a mass flow controller was directed through a bubbler before subsequently entering a 1-L glass bottle placed inside the temperature-controlled cabinet, which was set at either −10°C or 10°C depending on the experiment. The 1-L glass bottle acted as a condensing device, allowing excess water vapor to condense in the bottle and produce air leaving the condenser that was completely, or nearly, saturated with water vapor for that T_a_ (see results). Once humidified, air then flowed into a metabolic chamber. To prevent water vapor condensing and freezing in the tubes entering the condensing device and exiting the metabolic chambers, we inserted the tubes through copper pipes heated with a thermally controlled heating wire (Omega Engineering, Norwalk, USA). Excurrent air from a metabolic chamber first flowed through a water vapor analyzer (model RH-300, Sable Systems International, Las Vegas, NV USA) and subsequently scrubbed of CO_2_ and H_2_O prior to entering a FoxBox analyzer (Sable Systems International, Las Vegas, NV, USA,) for the measurement of 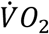. Air temperature within each metabolic chamber was measured with a type-T thermocouple connected to a Sable Systems thermocouple reader (model TC-2000, Las Vegas, NV, USA). Voltage outputs from all analyzers were digitized using a Universal Interface (model UI-2, Sable Systems International, Las Vegas, NV, USA) and recorded every 5 sec using a desktop computer with ExpeData software (v. 1.7.2, Sable Systems International, Las Vegas, NV, USA).

For each experimental sequence, birds were first exposed to dry, CO_2_-free air for 2 hours before being exposed to humidified air for another 2 hours. In both of the 2-hour sequences, the first hour was considered an acclimation period while the second hour was used in analyses. After the 4-hour experimental period, we removed the birds from their metabolic chambers and took a second measurement of T_b_ (within 1 min) and M_b_. Resting metabolic rate was calculated for each treatment by averaging the lowest, most stable 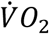 trace over a 10-min recording period. We used equation 10.1 from Lighton (2008) to calculate 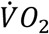 from incurrent and excurrent fractional gas concentrations.

Energy consumption was calculated assuming an RQ of 0.71 and a thermal equivalent of 19.8 kJ LO_2_^−1^ (Gessaman and Nagy, 1988). Final values were then converted to Watts. This protocol was approved by the Université du Québec à Rimouski Animal Care Committee.

### Data analysis

Our analyses first aimed at confirming that our experimental setup produced contrasting humid environments at the two experimental temperatures. To validate the humidity treatments, we ran the experimental setup without birds under dry and humid conditions. Measured WVP in an empty chamber was compared against the known saturation value for the average T_a_ recorded in the chamber. We used a three-way ANOVA to test for the effects of T_a_, humidity, and presence of bird in the chamber on WVP.

We used linear mixed effect models to test for the effects of T_a_ and humidity on RMR. We included Bird ID and measurement sequence as random effects in our models. We initially ran our analyses with mean M_b_ included as a covariate, where mean M_b_ represented the average of our M_b_ measurements taken before and after RMR runs. However, M_b_ was not significant (p > 0.05) and was removed from further analyses. All statistical analyses were performed in R (R Core Team, 2016 v. 3.2.3) using the *nlme* package (Pinheiro et al., 2016). Values presented are means with 95% confidence intervals (CI).

## RESULTS

### Experimental manipulation of humidity

Measurements of WVP in excurrent air from empty chambers confirmed that our experimental setup produced contrasting WVP (temperature x humidity x bird presence: F_1,58_ = 8.88, p = 0.004; Fig. 1). At below freezing T_a_, WVP in dry air was 0.03 ± 0.01 kPa and in humid air was 0.29 ± 0.02 kPa. At above freezing T_a_, WVP in dry air from empty chambers was 0.04 ± 0.02 kPa and in humid air was 1.03 ± 0.03 kPa.

**Fig. 1.**
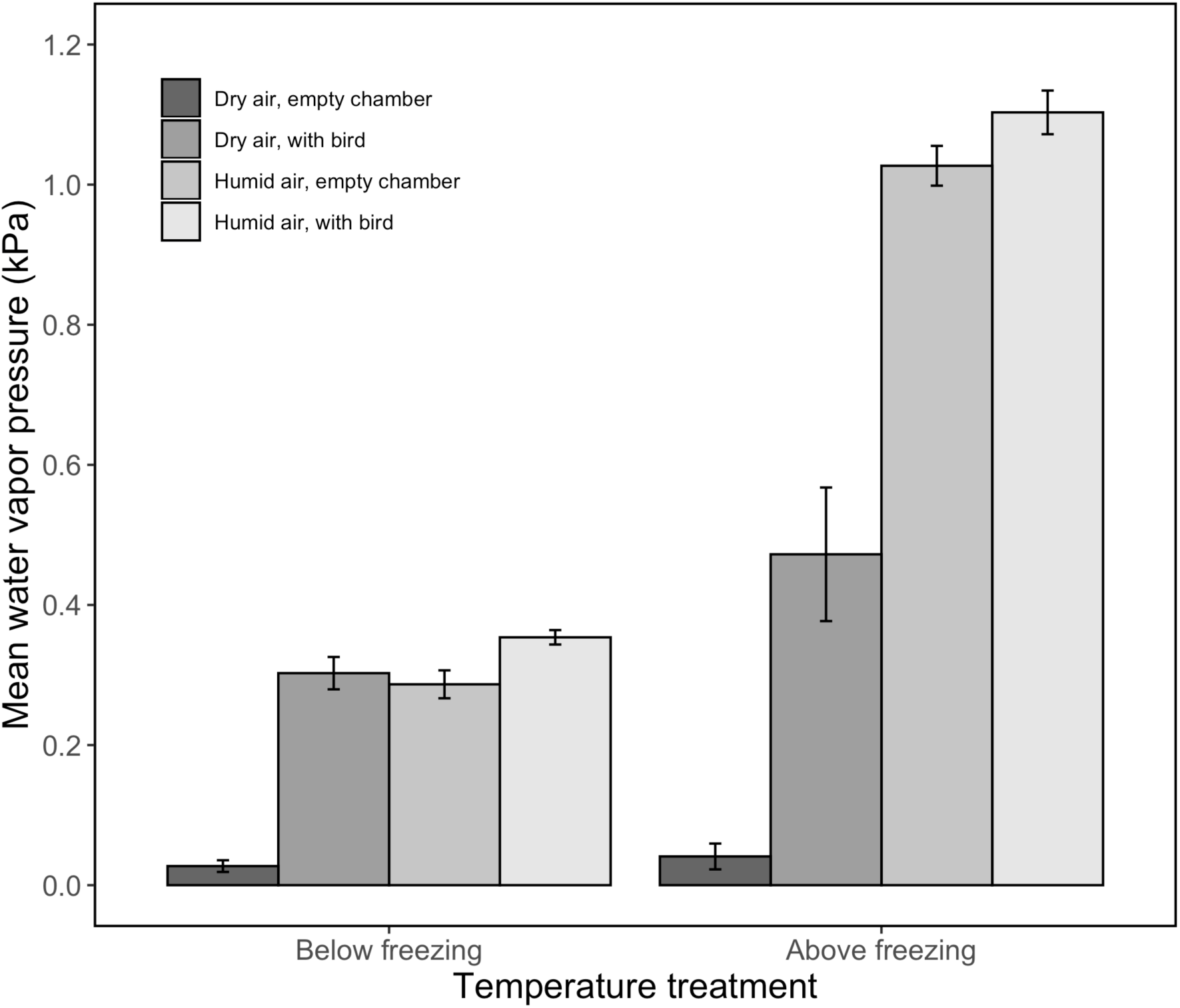
Mean water vapor pressure values recorded with flow-through respirometry for the two experimental temperatures and humidity treatments when a bird was either absent or present in a metabolic chamber. Error bars represent 95% confidence intervals.

Due to water vapor production through normal respiration, a bird’s presence in the chamber naturally raised humidity levels above those relative to empty chambers (Fig. 1).

For example, at above freezing T_a_, WVP in dry air increased to 0.47 ± 0.10 kPa (38.2% RH at T_a_ = 10.0°C) and in the humid treatment WVP raised to 1.10 ± 0.03 kPa (90.3% RH T_a_ = 9.9°C). At below freezing T_a_, birds exposed to dry air added enough water vapor so that humidity levels were similar to those in the humid treatment (Fig. 1).

Consequently, there was not a considerable difference in WVP between dry and humid treatments at below freezing T_a_ when chambers contained a bird (dry air: 0.30 ± 0.02 kPa [80.4% RH at T_a_ = −6.6°C]; humid air: 0.35 ± 0.01 kPa [96.7% RH at T_a_ = −7.0°C]).

### Resting metabolic rate

Temperature had a significant effect on RMR (F_1,8_ = 41.72, p < 0.001; Fig. 2). When all metabolic measurements were combined within a temperature treatment, average RMR was ~1.5 times higher at T_a_ below freezing (0.74 ± 0.06 W; mean ± SD T_a_ = −6.8 ± 0.6°C) than at T_a_ above freezing (0.51 ± 0.04 W; mean T_a_ = 9.9 ± 0.4°C). In contrast, humidity did not impact RMR at either T_a_ (temperature x humidity: F_1,17_ = 1.66, p = 0.22; Fig. 2).

**Fig. 2.**
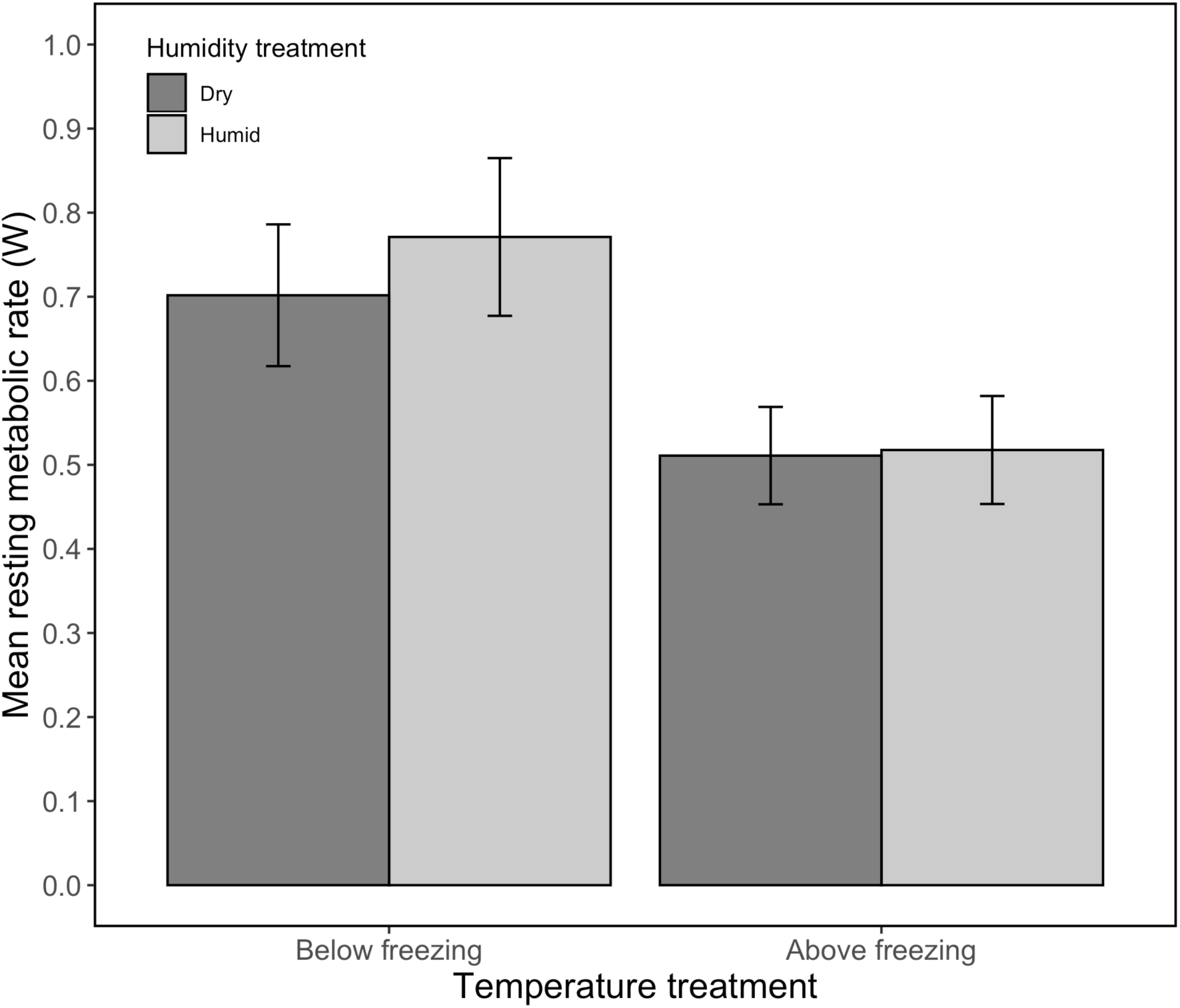
Mean resting metabolic rate of black-capped chickadees when exposed to dry or humid air at air temperatures (T_a_) below freezing (mean T_a_ for dry air = −6.6 °C and humid air = −7.0 °C) and above freezing (mean T_a_ for dry air = 10 °C and for humid air = 9.9 °C). Air temperature significantly influenced RMR while humidity had no effect. Error bars represent 95% confidence intervals.

At T_a_ below freezing, RMR under the dry treatment was 0.70 ± 0.08 W (mean T_a_ = −6.6 ± 0.8°C) and under the humid treatment it was 0.77 ± 0.09 W (mean T_a_ = −7.0 ± 0.5°C). At T_a_ above freezing, RMR under dry conditions was 0.51 ± 0.06 W (mean T_a_ = 10.0 ± 0.4°C) versus 0.52 ± 0.06 W (mean T_a_ = 9.9 ± 0.4°C) under humid conditions.

## DISCUSSION

In this study we assessed the scientific basis for the humid-cold belief widely held in northern populations by exposing Black-capped Chickadees to different combinations of T_a_ and humidity. We found that T_a_ had a significant effect on heat loss, manifested in Black-capped Chickadees increasing their RMR at colder T_a_ regardless of humidity level. The increased thermal gradient between T_b_ and T_a_ undoubtedly lead to increased heat loss to the environment which required increasing endogenous heat production to maintain an elevated T_b_ (Scholander et al., 1950). The increases in RMR observed here are comparable to inter-seasonal differences in metabolic rates reported for more southern Black-capped Chickadee populations (Cooper and Swanson, 1994; Sharbaugh, 2001), along with other northern wintering species under natural conditions (Hart, 1962; West, 1972; Olson et al., 2010; McKechnie et al., 2015). Liknes and Swanson (1996), for instance, found that basal metabolic rates were 49% higher in winter compared with summer in White-breasted Nuthatches (*Sitta carolinensis;* ~20 g) and Downy Woodpeckers (*Dryobates pubescens*; ~25g).

We predicted that at T_a_ below freezing a bird’s heat loss would not be influenced by humidity, whereas at T_a_ below the LCT, but still above freezing, birds would lose more heat under humid conditions resulting in a higher RMR. Our data, however, showed that humidity did not have a statistically significant influence on RMR in chickadees at either T_a_. Moreover, RMR actually exhibited a larger difference between humidity groups at T_a_ below freezing, contrary to our prediction. However, the effect size was very small, especially when compared against the effect of T_a_ on RMR (i.e., a difference of 0.07 W among humid treatments within T_a_ below freezing *versus* a 0.23 W difference among the two T_a_ treatments). A likely reason for the absence of a considerable humid effect at T_a_ below freezing reflects the fact that the absolute amount of water that the air can hold at this temperature is extremely low. For example, saturated air at −10°C contains ≈24% of the water contained at 10°C (2.3 mgH_2_O L^−1^ of air *versus* 9.4 mgH_2_O L^−1^ of air, respectively). Thus, the amount of water vapor in the air at air temperatures below freezing is so low that any potential increases in heat conduction away from the body is certainly negligible.

It is important to note here that because of a bird’s contribution in water vapor through respiration, WVP values in the dry treatment were similar to those in the humid treatment at T_a_ below freezing. Consequently, any humidity effect was presumably negated, and the small differences observed in RMR between the humid and dry treatments cannot unequivocally be ascribed to humidity level. However, because we observed no difference between RMR values between humidity treatments at T_a_ above freezing, we are confident that this has no substantial impact on our final conclusions.

So, if decreasing air temperatures hold less water vapor, likely making any possible increases in heat transfer through conduction negligible, why does the humid-cold perception persist? A likely answer stems from the concept of environmental, or operative, temperature (Bakken et al., 1976; Robinson et al., 1976). The thermal environment that an organism experiences is often derived from the interaction among several parameters, namely, T_a_, wind, solar and thermal radiation, and humidity. Solar radiation, in particular, can significantly contribute to the operative temperature experienced by an organism (Fortin et al., 2000; O’Connor et al., 2018). It is noteworthy that the Canadian cities regularly reporting to feel warmer during winter periods because of dry-cold conditions (e.g., Calgary and Edmonton) are some of the most sunlit areas, whereas those known for humid-cold (e.g., Toronto and Vancouver) are some of the cloudiest regions during winter (Fig. 3). Hence, we agree with Crow’s (1988) suggestion that the presence (or absence) of solar radiation, and not water vapor, is a more likely culprit behind the humid-cold myth.

**Fig. 3.**
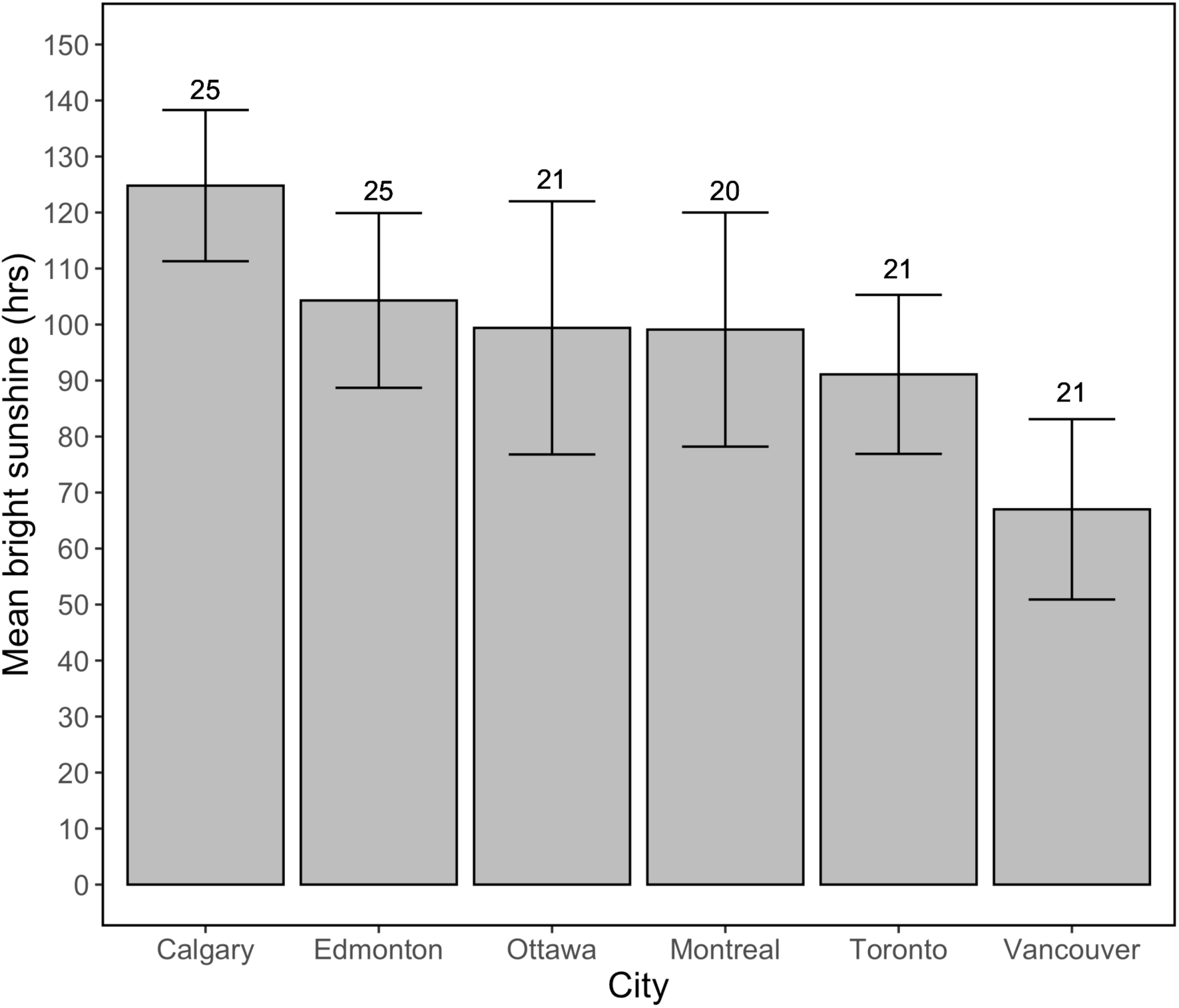
Mean hours of bright sunshine for several Canadian cities spanning the period from November to February between 1981-2010. Error bars represent standard deviation and the numbers above bars represent the total number of years over which the data were collected. Data are from the Canadian Climate Normals 1981-2010 (http://climate.weather.gc.ca/climate_normals/index_e.html?#1981).

## List of symbols and abbreviations

M_b_: Body mass
LCT: Lower critical temperature
RH: Relative humidity
RMR: Resting metabolic rate
T_a_: air temperature
T_b_: Body temperature
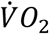: Oxygen consumption;
WVP: water vapor pressure

## Acknowledgements

We greatly thank Justine Drolet for substantial help with the collection and organization of data. We thank Lyette Régimbald for her help with the captive birds and Alain Caron and Vincent Gauthray-Guyénet for their statistical advice. We are also grateful to Alexander R. Gerson and Scott McWilliams for their suggestions on interpreting results. We also thank Michal Wojciechowski for his advice on using condensing devices in our experimental setup.

## Author contributions

Conceptualization: F.V.; Formal analysis: F.V., R.S.O.; Investigation: F.V.; Writing – original draft: F.V.; Writing – review & editing: R.S.O., F.V.; Visualization: R.S.O.

